# Lipid-driven SRC self-association modulates its transformation capacity

**DOI:** 10.1101/2024.11.12.623151

**Authors:** Irrem-Laareb Mohammad, Marina I. Giannotti, Elise Fourgous, Yvan Boublik, Alejandro Fernándz, Anabel-Lise Le Roux, Audrey Sirvent, Marta Taulés, Serge Roche, Miquel Pons

## Abstract

Src tyrosine kinase is anchored to the plasma membrane, however, the contribution of membrane lipids to its regulation remains elusive. Here we report Src self-association through a lysine cluster in the Src SH4 region mediated by lipids in human cells and in vitro. Mutation of the lysine cluster to arginine modulates Src self-association and its transforming function in human cells. This mechanism may also apply to other membrane-associated signaling proteins that have similar lysine clusters in their unstructured regions.

Lipid-anchored micron-sized condensates of full length Src are formed in supported homogeneous lipid bilayers (i.e independently of lipid phase separation). Condensates are also formed by the purified Src N-terminal regulatory element, including the myristoylated SH4 domain, the intrinsically disordered Unique domain and the globular SH3 domain. The isolated SH4 domain forms small protein-lipid clusters, but not micron-size condensates.

Our findings reveal lipid-mediated kinase self-association as an additional mechanism for Src regulation.

**Significance Statement:** Condensate formation, often associated to intrinsically disordered protein regions is emerging as a widespread regulatory mechanism. Protein phase separation has been linked to the regulation of kinases recruited to condensates formed by other proteins. Here we show i) that the N-terminal disordered region of Src kinase drives lipid-mediated self-association to form condensates on the cell surface, ii) a lysine cluster in the N-terminal SH4 domain is essential for this process, iii) mutation of these lysine residues to arginine increases self-association in vitro and in vivo, and iv) self-association of Src results in enhanced transforming capacity in human cells. Similar lysine clusters are present in juxtamembrane regions suggest a general mechanism for modulating protein-protein interactions by lipids.

The article shows the formation of condensates by Src on supported lipid membranes mediated by lipids, dependent of a conserved lysine cluster in its SH4 domain, and modulating Src’s transforming capacity.

- Full length Src and truncated variants including its myristoylated SH4 domain self-associate on the surface of homogeneous lipid bilayers.
- The interaction of a lysine cluster in SH4 domain with lipids modulates Src self-association
- Atomic force microscopy shows the formation of micron-size condensates and smaller clusters on the surface of supported lipid bilayers, depending on the lipid composition or the presence of mutations in the lysine cluster
- Src self-association is confirmed in human cells and the Src transforming function of Src in human cells is modulated by lipid-mediated self-association

## Introduction

The non-receptor Src family (SFK) protein tyrosine kinases (TKs) are key regulators of membrane signaling that controls cell growth and adhesion.^1^ Src, originally identified as an oncogene, is the prototypical SFK. It shares with other SFKs a common modular structure consisting of an SH3, SH2 and a kinase domain (SH1) followed by a short C-terminal tail (scheme 1). ^2^ Additionally, Src contains an N-terminal unstructured region composed of a short myristoylated SH4 domain and a unique domain (UD) with unclear function.^3^ Deregulation of Src is frequently observed in human cancers and has been linked to tumor progression. ^4,5^ However, SRC oncogenic mutations are rarely detected in human cancer, suggesting the existence of important non-genetic mechanisms that drive Src kinase over-activation and cell transformation.

Crystallographic studies have identified a key conformational mechanism for regulation of Src catalysis mediated by SH2- and SH3-dependent intramolecular interactions.^6,7^ However, this core regulatory mechanism alone may not explain the full diversity of Src signaling and functions.

Myristoylated SH4 is essential for Src anchoring to the inner layer of the cell membrane and transforming, but kinase regulation in this lipid context, remains elusive. Cell membrane partition through cholesterol-enriched lipid rafts was reported to induce Src clustering with their receptors, enabling signaling^8–10^. Src dimerization involving its disordered N-terminal region has been reported in mammalian cells and related to the enhanced phosphorylation of selected substrates^11^. However, in vitro studies have demonstrated that self-association of Src N-terminal region does not require lipid-separated phases^12^. Recent reports^13^ and reviews^14^ highlight the possible role of liquid-liquid protein phase separation in the regulation of kinases. Intrinsically disordered regions are often involved in the formation of phase separated liquid protein condensates^15–17^. The Src N-terminal intrinsically disordered region modulates its membrane anchoring, enabling membrane substrate phosphorylation and oncogenic signaling.^18^ Mechanistically, the disordered domains make a fuzzy complex condensed around the SH3 domain, while retaining the extensive dynamics typical of intrinsically disordered regions,^19,20^ and modulate substrate selection and signaling. ^21^ When anchored to membranes, the myristoylated disordered region enables *in vitro* self-association,^12,22^ although the detailed mechanism is currently missing. Overall, these data point to an additional layer of Src regulation implicating its unstructured region and membrane lipids. Here, we explored the mechanism for crosstalk between this unstructured region and membrane lipids by focusing on lipid mediated Src self-association (SA).

To this end we used the following strategy. First, using surface plasmon resonance to detect SA, we searched for the SH4 residues that were essential for Src SA in the context of myristoylated SH4-UD-SH3 (named Src N-terminal Regulatory Element, SNRE) (Figure 1A). Next, to explore the nature of Src SA and to confirm that it occurs *in vitro* also in the context of larger constructs, including full length Src, we used atomic force microscopy. We used neutral and anionic supported lipid bilayers that do not form lipid-separated phases to demonstrate that the capacity of Src to SA is intrinsic and independent of lipid phase separation. Having identified the essential role of a SH4 lysine cluster in Src SA and demonstrated that replacement of these residues by arginine enhances SA, we translated these *in vitro* results to human cells, demonstrating that the lysine to arginine mutations in the SH4 domain also enhance Src SA *in vivo* and modulates Src transforming capacity.

**Figure 1.**
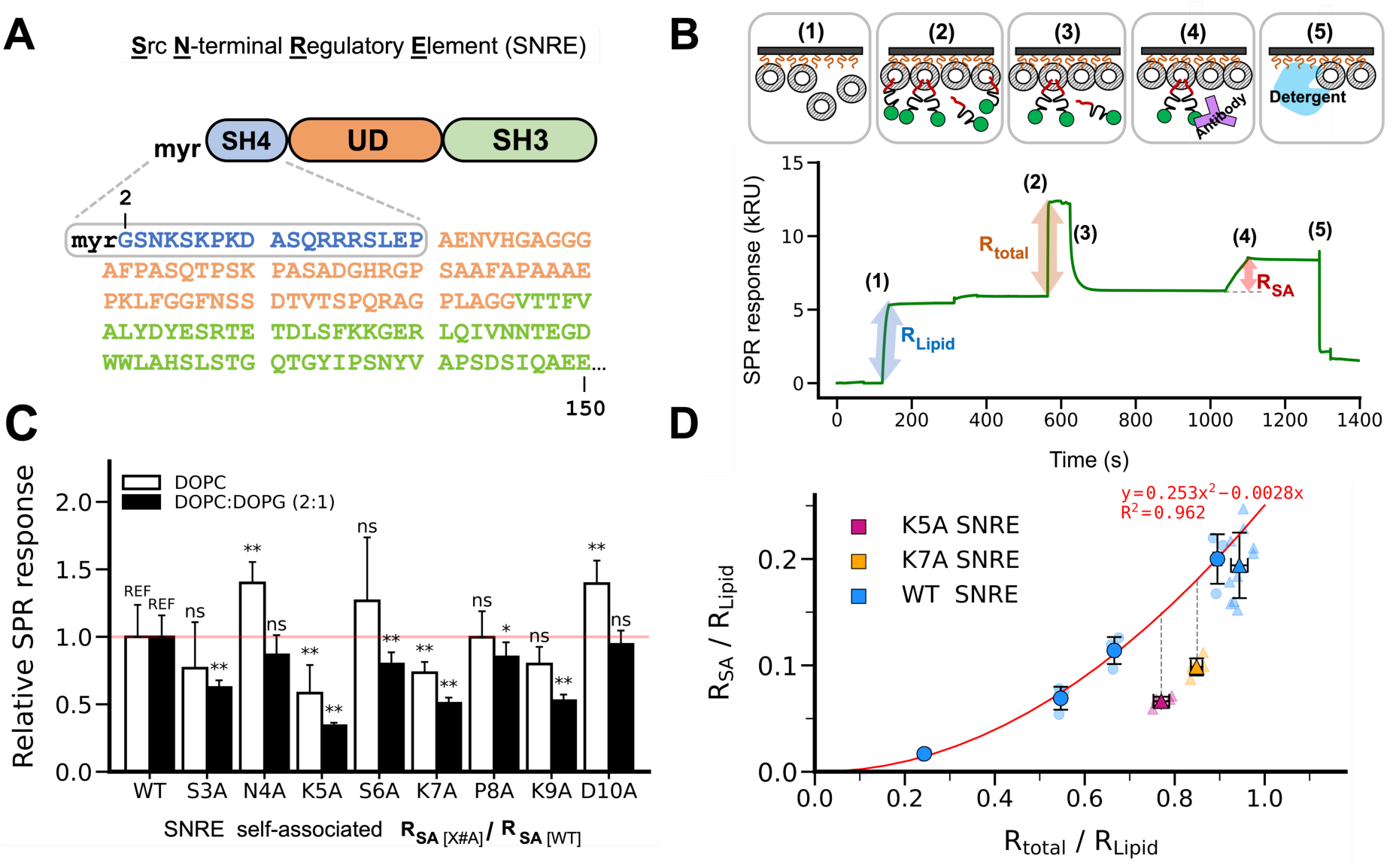
A lysine cluster in the SH4 domain participates in lipid-mediated SNRE dimerization. (A). Sequence of the Src N-terminal regulatory element. The SH4 and UD are intrinsically disordered. (B) Experimental protocol used to quantify self-association by Surface Plasmon Resonance (SPR). Liposomes were immobilized using a phytosphyngosine derivatized matrix. Total bound protein was quantified by the direct SPR response. Following washing, only persistently bound species arising from self-association were retained and quantified by the response to an antibody not interfering with membrane binding. (C) Self-association quantification of SNRE mutants relative to the WT form using pure zwitterionic DOPC (open bars) or a mixture of DOPC with negatively charged DOPG (filled bars). Proteins were injected at 20 μM. Data expressed as mean ± SD, n = 4 (DOPC) n=5 (DOPC: DOPG). Significant differences are indicated with asterisks (Student’s t-test: *p < 0.1; **p < 0.05; ns: not significant; REF: reference). D) The effect of K5 and K7 mutations on SA is not due to reduced membrane affinity of the monomers. WT SNRE was injected at a bulk concentration of 3, 6, 10 and 20 μM whereas the K#A mutants were injected at 20 μM. Data used originates from two different sensor chips indicated by the type of marker (circle or triangle). Data expressed as mean ± SD, n = 3 (circles) n ≥ 5 (triangles).

## Results

### The SH4 K5, K7 and K9 modulate lipid-mediated SNRE self-association

In a prior study, we designed an *in vitro* assay, shown schematically in Figure 1B, to assess Src binding kinetics onto membrane lipids using Surface Plasmon Resonance (SPR).^22^ For this, we used immobilized lipid bilayers (SLB) with purified SNRE from *E. coli* co-expressing yeast N-myristoyl Transferase to induce protein myristoylation. Our SPR study revealed a major lipid bound monomeric SNRE population and a minor form that was persistently bound to the lipid membrane. Kinetic studies using SPR,^22^ and single molecule fluorescence^12^ indicated that the persistently bound form involves at least two SNRE molecules. Here we investigated the underlying mechanism by focusing on the contribution of the SH4 region by an alanine-scan targeting residues 3 to 10. We focused in the initial region of the domain based on previous studies that showed an effect of deletion of the first 10 residues of the SH4 domain in the compaction of the fuzzy complex with SH3^19^ and the lipid membrane distribution and clustering.^23^

We first used SLB composed of neutral zwitterionic lipid 1,2-dioleoyl-sn-glycero-3-phospho choline (DOPC) in liquid disordered (L_d_) phase. The relative SPR responses of the individual mutants with respect to wild type (WT) SNRE showed that replacing lysine residues in positions 5 or 7 by alanine (K5A and K7A, respectively) reduces SA by around 30-40% (Figure 1C), while replacement of the negatively charged aspartic acid by alanine (D10A) increases it. An SA increase is also observed in the N4A mutant in neutral lipids. We next assessed the contribution of anionic lipids. For this, we used a 2:1 mixture of DOPC and 1,2-dioleoyl-sn-glycero-3-phospho(1’-rac glycerol) (DOPG), which does not form separate phases in our conditions,^24^ avoiding the possible complications caused by changes in lipid-phases induced by the protein, while still allowing to test the relative effects with zwitterionic lipids. Interestingly, inhibition of SA induced by K5A and K7A mutations was enhanced by 10% and an inhibition was observed also in the K9A mutant, thus implicating an SH4 lysine cluster and negative charged lipids for SNRE SA (Figure 1C). Mutation of serine 3 (S3A) also decreases SA on anionic bilayers. This residue is located between the lysine cluster and the myristoylation site. We suggest that it influences SA by allowing the positioning of the positively charged residues when the myristoyl group is inserted in the lipid bilayer.

### K5A and K7A mutations hinder SNRE self-association independently of their effect on Src membrane affinity

Because SH4 K#A mutations may also affect the lipid binding affinity of individual Src molecules and, therefore, the actual concentration of Src on the membrane surface, we compared SNRE SA capacity at the same total density of bound proteins. These experimental conditions were obtained by varying the bulk concentration of injected SNRE variants (3 to 20 μM). In conditions of same bound SNRE density, SA formation was still reduced by 40% for K7A and 50% for K5A with respect to WT (Figure 1D). Therefore, the observed decrease in SA population in the K#A mutants cannot be explained only by a reduction of the protein density on the lipid surface. Interestingly, the steady-state response of the K5A and K7A mutants was not the same, despite having the same overall charge, suggesting that the position of the basic residues is important for primary binding and SA. We conclude that the lysine cluster in the SH4 is driving SNRE SA in the presence of lipids, beyond its electrostatic contribution to the binding to negatively charged lipid membranes and despite the expected electrostatic repulsion between adjacent Src molecules.

### Src forms membrane anchored condensates observable by atomic force microscopy on supported lipid bilayers

The requirement of a cluster of positive charged residues for SA of Src on the membrane surface suggests that the negatively charged phosphate group of phospholipids could mediate the interaction between Src molecules, however, SPR does not provide information on the nature of the species formed by SA. Therefore, we studied Src SA by atomic force microscopy (AFM). Addition of SNRE to an SLB made of pure DOPC led to the formation of micron-size condensates with a smooth surface about 1.5 nm thinner than the surrounding lipid bilayer (Figure 2A). Micron-size condensates were also observed with myristoylated full length Src (Figures 2B), although the appearance of the condensates formed by full length Src was different from those formed by the truncated protein. First, the thickness of full length Src condensates is similar or exceeds that of the surrounding bilayer, which is compatible with the much larger size of the protein (150 vs 536 residues for SNRE and full-length Src respectively). The second difference is that the border region of the full length Src condensate has a complex structure in contrast with the plain border of the condensates formed by SNRE.

**Figure 2.**
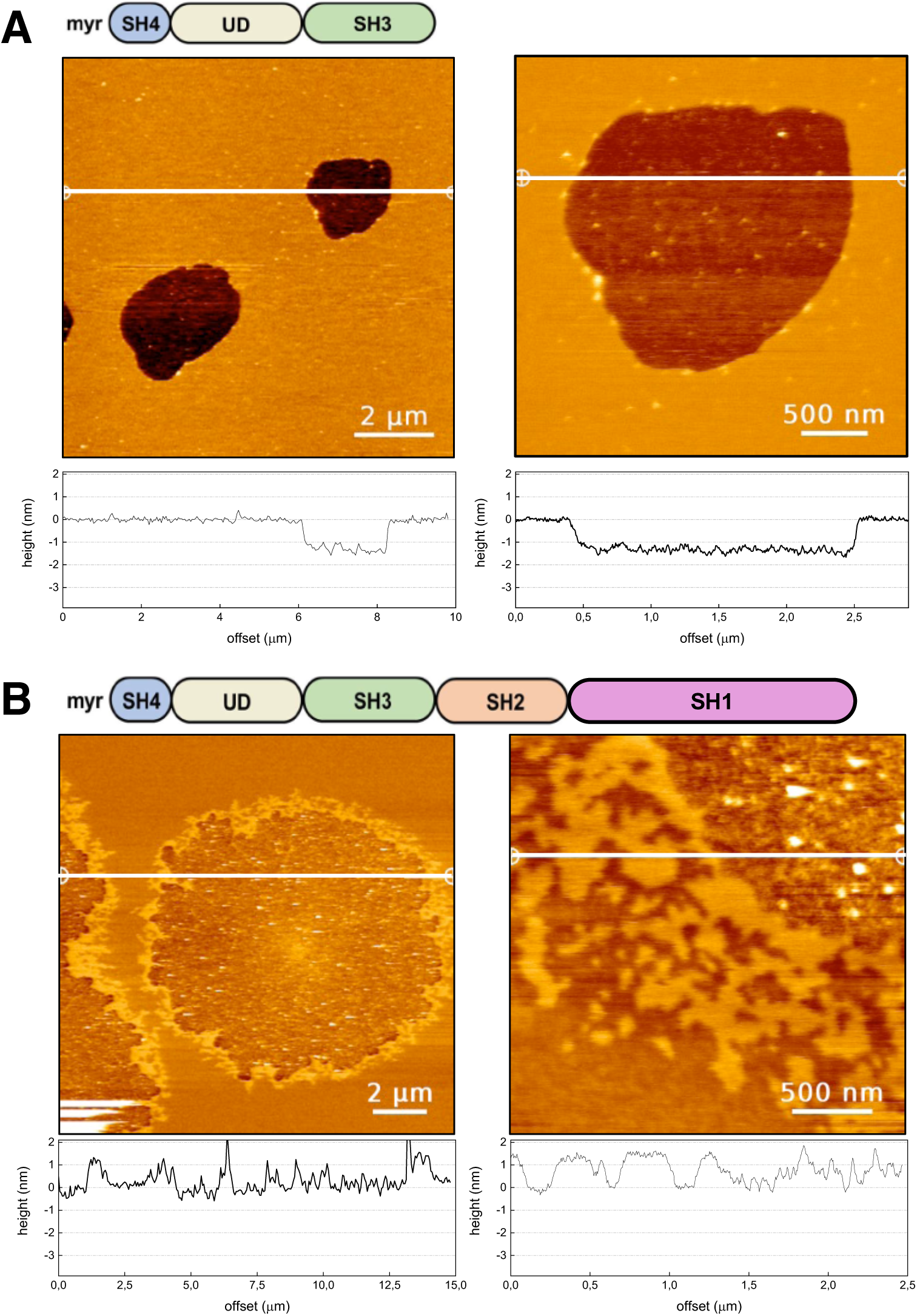
Condensate formation by SNRE and full length Src on supported pure DOPC lipid bilayers. Topographical AFM images of SNRE (A) and myristoylated full length Src on mica-supported lipid bilayers (SLB) made of pure DOPC. Cross sections along the indicated lines are shown underneath each image. The protein was added to the buffer solution covering the SLB, to a final concentration of 100 nM, and left to interact for 4 min before imaging. Images were acquired in working buffer (10 mM NaH_2_PO_4_/Na_2_HPO_4_, 150 mM NaCl, pH 7.5)

We next analyzed lipid mediated self-association in SLBs made of DOPC:DOPG 4:1 that form homogeneous L_d_ membranes. Indeed, homogeneous lipid bilayers were observed in the absence of protein (Figure 3A). Micron-size condensates were also observed upon addition of SNRE (Figure 3B and 3D).

**Figure 3.**
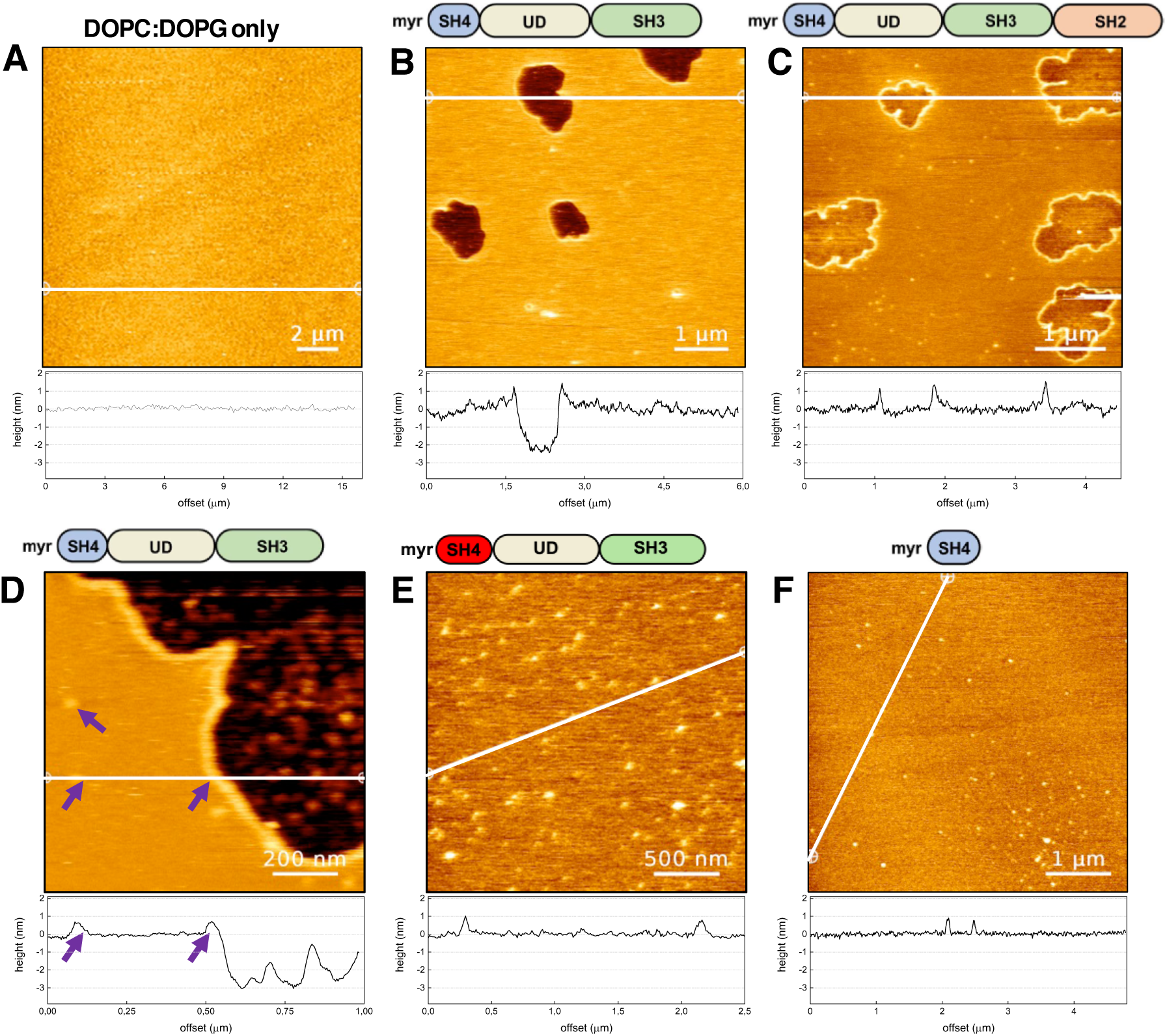
Condensate formation by truncated Src forms in negatively charged DOPC:DOPG 4:1 supported lipid bilayers. (A) A homogeneous bilayer formed in the absence of protein confirms that DOPC:DOPG mixtures do not form separated lipid phases. (B) Condensates formed by SNRE. (C) Condensates formed by a larger construct including the SH2 domain. (D) Coexistence of micron- and nm-sized condensates formed by SNRE. Notice the similar height of the nm-sized clusters and the wall surrounding the micron-sized condensates. (E) Only nm-sized clusters are formed in the 3R mutant of SNRE. (F) Only nm-sized clusters are formed by the isolated myristoylated SH4 domain.

The presence of negatively charged lipids caused some differences in the appearance of the SNRE condensates, as compared to those observed with a single, zwitterionic, lipid species (DOPC). First, the thickness of the condensates was not homogeneous with a height profile changing abruptly between 2 and 3 nm below than the surrounding lipid bilayer. Second, the condensates formed in the presence of negatively charged lipids were surrounded by a “wall” protruding around 0.5 to 1 nm above the lipid surface (Figure 3D), that was not present in the absence of DOPG. In addition to the micron-sized condensates, many small (nm-size) clusters with a heigh of 0.5 nm comparable to that of the wall surrounding the condensates were observed only in the presence of negatively charged lipids. These features suggest the preferential interaction of the protein with negatively charged lipids, leading to the formation of smaller lipid-protein clusters that can either remain isolated, located on the border of the larger condensates forming a wall, or become incorporated in the condensates, explaining the rougher surface of the condensates formed in the presence of negatively charged lipids, as compared to those formed in pure DOPC.

Similar features were observed with a larger construct incorporating the SH2 domain (Figure 3C), except for the thickness of the condensate, that is intermediate between that observed for the myr-SH4-UD-SH3 construct and full length Src in DOPC. Unfortunately, full length Src caused the detachment of the lipid from the mica surface and could not be studied in DOPC:DOPG bilayers.

### Formation of small lipid-protein clusters is mediated by the SH4 domain but micron size condensates require the presence of the Unique domain

If protein-lipid clusters are mediated by the lysine-cluster, as suggested by the SPR experiments, replacement of lysine by arginine that is known to have a stronger interaction with phosphate groups^25^ may affect the balance between discrete clusters and large condensates. Thus, we tested an SNRE mutant in which K5, K7 and K9 had been replaced by arginine residues (3R mutant). Indeed, 3R SNRE formed many small clusters but no micron-size condensates (Figure 3E).

Since the formation of condensates is often associated to intrinsically disordered regions, we tested just the myristoylated SH4 domain, lacking the intrinsically disordered Unique domain (UD). Myristoylated SH4 only forms discrete clusters (Figure 3F), confirming that while Src self-association involves the SH4 domain, formation of large condensates requires additional regions.

Self-association, the role of the lysine cluster, and the enhanced self-association caused by the 3R mutation was confirmed in the isolated SH4 domain by SPR (Supplementary figure S1).

Overall, the AFM and SPR experiments support the hypothesis of a lipid mediated SNRE self-association mechanism and the importance of SH4 lysine cluster and anionic lipids in this molecular process.

### The SH4 lysine cluster regulates Src self-association in human cells

We next sought experimental evidence for Src SA in human cells. For this, full length Src constructs tagged with a Myc or a Flag sequence at the C terminus were co-transfected in HEK293T cells. We first confirmed by western blot analysis of the expressed proteins and the cell’s phosphoprotein profile, that adding these tags at the regulatory tail did not affect Src expression or activity of WT Src (Figure S2). Based on the results obtained by SPR and AFM showing enhanced SA by the 3R mutants and the fact changing lysine to arginine does not affect in vivo myristoylation,^26^ we compared SA of WT and 3R full length Src in HEK293T cells. Confocal microscopy confirmed similar subcellular distribution between Src and Src3R, notably at the plasma membrane (Figure S3). Using the tagged proteins, Src SA was clearly detected by co-immunoprecipitation. Importantly, arginine replacement of the SH4 lysine cluster (Src3R) increased Src SA by 50% (Figure 4A). Src SA in cells was independently confirmed by proximity ligation assay (PLA) that does not depend on immunoprecipitation but detects protein pairs separated by a distance lower than ca. 40 nm using immunofluorescence detection. Our approach was first validated by showing Src association with the pseudo-kinase PEAK2, a well-established Src substrate and interactor (Figure S4).^27^ Src3R showed a 100% increase of PLA-detected SA as compared to Src WT (Figure 4B and S4). Overall, these data agree with our *in vitro* findings and confirm that Src SA also occurs in vivo, supporting our model of lipid-driven SA of membrane anchored-Src.

**Figure 4.**
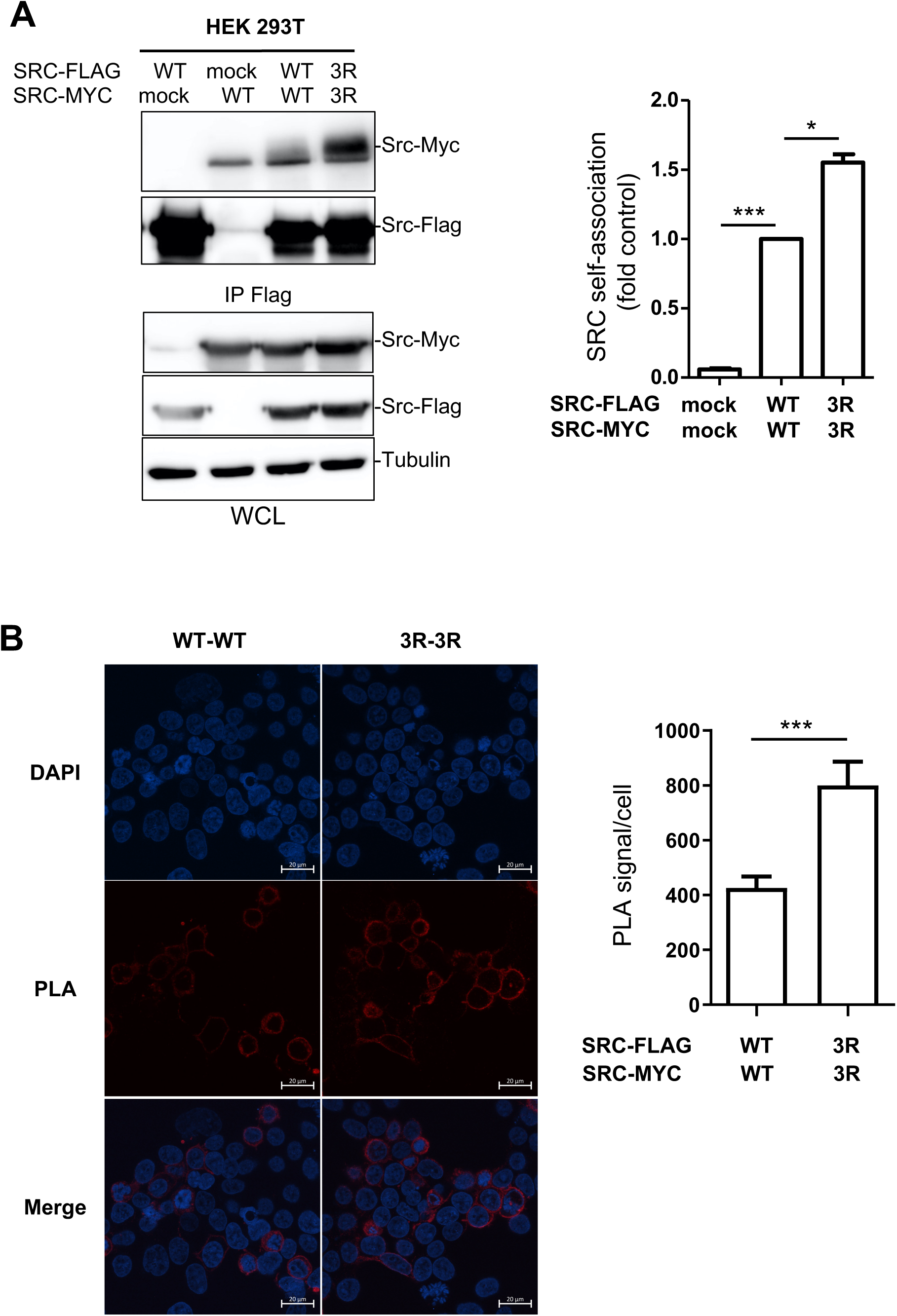
Mutation of lysine to arginine in the SH4 lysine cluster enhances Src self-association in human cells. (A): HEK293T cells were transfected with the indicated Src constructs. Src-Flag proteins were immunoprecipitated (IP) from cell lysates and immunoblotted with the indicated antibodies. Immunoblots of whole-cell lysates were also performed as indicated. A representative example and the relative quantification of co-immunoprecipitation is shown. (B): Proximity ligation assay (PLA) from cells transfected with indicated construct was used to prove the proximity of Src variants in intact cells. A representative example and the relative quantification of PLA signal/cell, as assessed by confocal microscopy, is shown as the mean ± SEM; n=3-5; ns: p>0.05; *p<0.05; **p<0.01; ***p<0.001; Student’s t test. Control experiments are presented in supplementary figures S2 (expression and activity of tagged WT Src), S3 (colocalization of WT and 3R Src), and S4 (positive PLA control and whole-cell lysates).

### Src SA increases oncogenic signaling in human cancer cells

Finally, we evaluated the functional relevance of Src SA, by assessing the impact of Src3R mutants on cell signaling. Src3R led to a 50% increase in cellular protein tyrosine phosphorylation consistent with an increased kinase activation (as assessed on pY419 levels) (Figure 5A). Finally, we evaluated the effect of Src3R on Src transforming functions. We previously reported that Src overexpression enhanced transforming properties of colorectal cancer cells, including the HCT116 cell-line,^28^ because of inhibition of key control mechanisms.^29^ Using this model, we observed here that Src3R was more oncogenic than WT Src, as assessed on anchorage-independent growth and cell migration in Boyden chamber (Figure 5B). We thus concluded that Src lipid-driven SA modulates Src signaling and oncogenic function.

**Figure 5.**
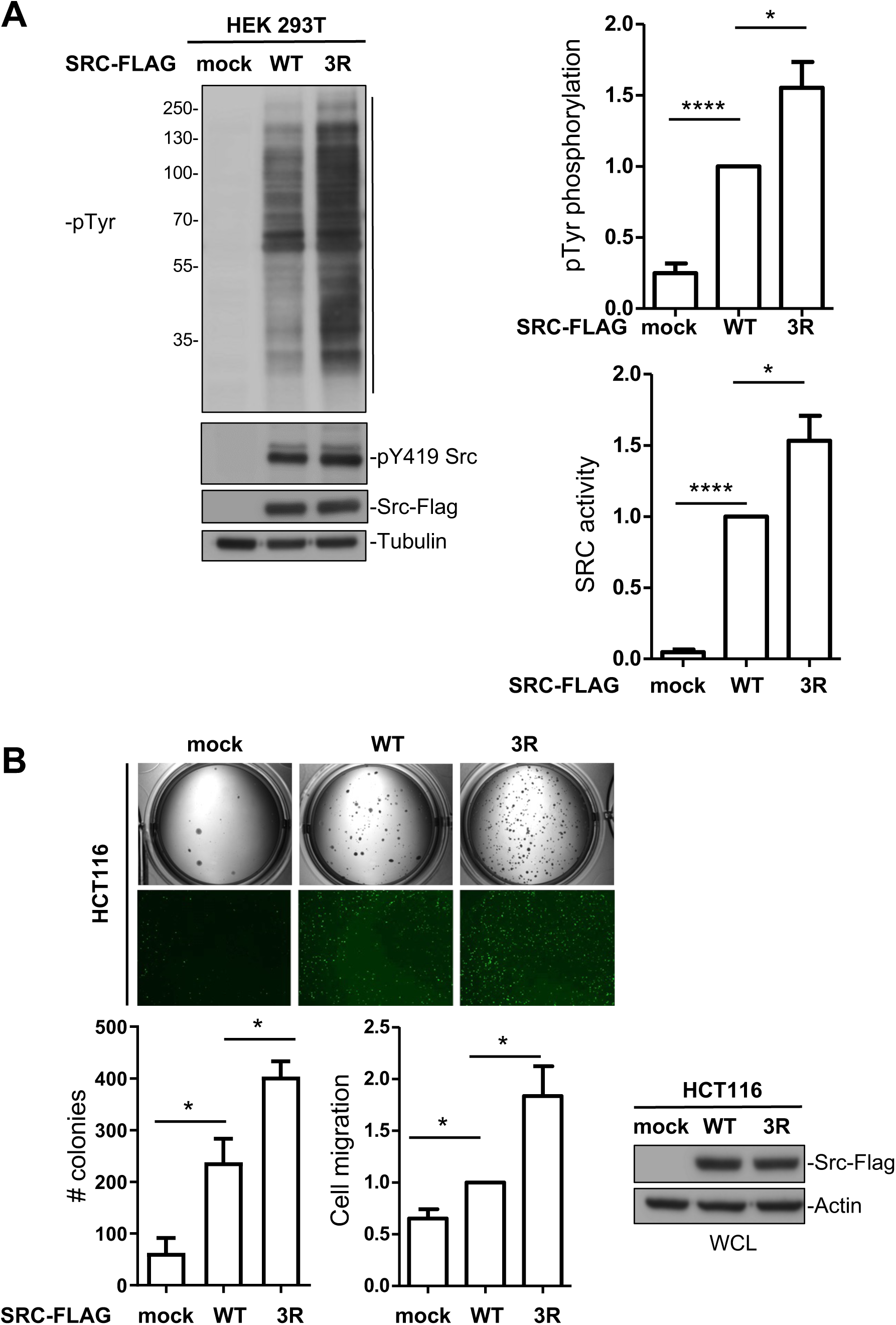
Src self-association increases oncogenic signaling in human cancer cells. (A): phospho-signaling of indicated c-Src variant transfected in HEK293T cells. A representative example and quantification of c-Src induced protein tyrosine phosphorylation (pTyr) and Src activity (pY419 Src) assessed by immunoblotting. (B): transforming activity of WT and 3R c-Src retrovirally transduced in human colorectal cancer HCT116 cell-line. An example (top panels) and quantification of c-Src induced colonies formation in soft agar and migration in Boyden chambers. The level of transduced c-Src is also shown (bottom right). Is shown the mean ± SEM; n=3-5; ns: p>0.05; *p<0.05; **p<0.01; ***p<0.001; Student’s t test.

## Discussion

### A novel mechanism of Src regulation by membrane lipids

Here we provide further evidence to support a mechanism for Src regulation through the interaction with membrane lipids. We propose a model where a cluster of positively charged lysine residues in the SH4 domain and negatively charged lipids drive Src SA.

The importance of the cluster of lysine residues in the SH4 domain of Src was already recognized in the early work by the groups of Bishop, Varmus, Resh and McLaughlin.^26,30,31^ They established its role for in vivo myristoylation by N-myristoyltransferase and, more importantly, the importance of their electrostatic interaction with negatively charged lipids to enhance the anchoring of individual Src molecules to the membrane. However, the role of the SH4 lysine cluster as driver for Src self-association was previously unknown. We now show that the lysine cluster in the SH4 domains promotes lipid-mediated Src self-association on the membrane surface. We have also demonstrated that mutation of the lysine residues in the cluster to arginine results in much enhanced self-association in vitro and in vivo and leads to an increased transforming capacity in HCT116 colorectal cancer cells.

Previous results from our group had shown that the SH4 domain also participates in the formation of a fuzzy intramolecular complex, involving the Unique and SH3 domains. Fuzzy complexes are one of the consequences of multivalency, the coexistence of alternative weak interactions that, collectively lead to a macroscopically strong effect. Another extensively found manifestation of multivalency is the formation of separate phases or condensates, often associated to disordered regions and including a variety of components. Using atomic force microscopy, we have observed the formation of micron-sized condensates by full length Src and various truncated forms in supported lipid membranes. Condensates are formed by lipid-mediated protein-protein interactions. In the presence of negatively charged lipids, smaller protein-lipid clusters are formed coexisting with the large condensates. Presumably, the small clusters result from the recruitment of negatively charged lipids by a small number of protein molecules, while the larger condensates result from weaker, more delocalized interactions, and possibly direct protein-protein interactions away from the membrane surface. Interestingly the isolated myristoylated SH4 domain and a longer construct with the 3R mutation form only small clusters, suggesting that lipid-mediated self-association is the first step of a more complex mechanism involving other Src regions.

### Lysine clusters are abundant in juxtamembrane regions: a possible general mechanism for lipid-driven protein-protein interactions

Because alternating lysine clusters are observed in other SFKs,^32^ including Fyn (^7^KDKEATK^13^), Yes (^5^KSKENK^10^) and Lyn (^5^KSKGKD^10^), we predict that a similar lipid-mediated regulatory mechanism may apply to these SFK members. Interestingly, membrane anchorage of these SFK is already stabilized by protein palmitoylation; therefore, this conserved lysine cluster would contribute to association with other SFK kinases (homo or mixed dimers), enabling SFK signaling specificity and diversity. While the abundance of positively charged residues in the juxtamembrane region of integral membrane proteins has been long recognized, our results suggest that, as in the case of Src, clusters of basic residues may be a general motif involved in lipid-mediated protein-protein interactions and membrane signaling. Consistent with this idea, a similar positive cluster motif is present in membrane-anchored K-Ras isoform 2B (^180^KSKTK^184^) forming nanoclusters.^33^ Lipid-driven protein-protein interactions offer yet an additional regulatory layer exploiting the large variability in lipid composition of different types of cells or even among cells of the same type in different metabolic states. The concept of lipotypes has been suggested to describe stable lipid configurations, associated to specific transcriptional programs and cell phenotypes.^34^ The identification of lipid-driven association mechanism in Src, may contribute to Src oncogenic induction in tumor cells harboring aberrant lipid metabolism with obvious implications in cancer therapy.

## Acknowledgements

We acknowledge the assistance of Roger Martínez in the preparation of some mutants, Javier Carvajal in some SPR measurements, Daniel Bouvard (CRBM, Montpellier) for confocal microscopy and Montpellier RIO Imaging for imaging analysis. This work has been funded by the Spanish Agencia Estatal de Investigación (PID2019-104914RB-I00, PDC2021-121629-I00, financed by European Union Next Generation Funds, and PID2022-140459OB-I00, cofunded by MICIU/AEI/10.13039/501100011033/ and FEDER A way of making Europe), la Ligue Nationale Contre le Cancer (LNCC), Montpellier SIRIC Grant «INCa-DGOSInserm 6045», the Agence Nationale de Recherche (FUZZY-SRC ANR-21-CE13-0011), and and CIBER (Consorcio Centro de Investigación Biomédica en Red, CB06/01/008.

## Authors contributions

I-L M: Experiment design, experiment execution and analysis of SPR data. M.I.G: AFM experiments and analysis. E.F.: Biochemical and cell biology experiments. Y.B.: Biochemical and cell biology experiments. A.F.: Preparation and characterization of myristoylated proteins. A-L. LR: SPR and AFM experiments. A.S.: PLA studies. M.T.: SPR methodology. S.R.: Design and supervision of biochemistry and cell biology experiments. Funding acquisition, Manuscript writing. M.P.: Conceptual design. General supervision. Funding acquisition. Manuscript writing.

## Data availability

Original data created for the study will be available in the Zenodo repository upon publication.

## Figure legends

**Figure S1.**
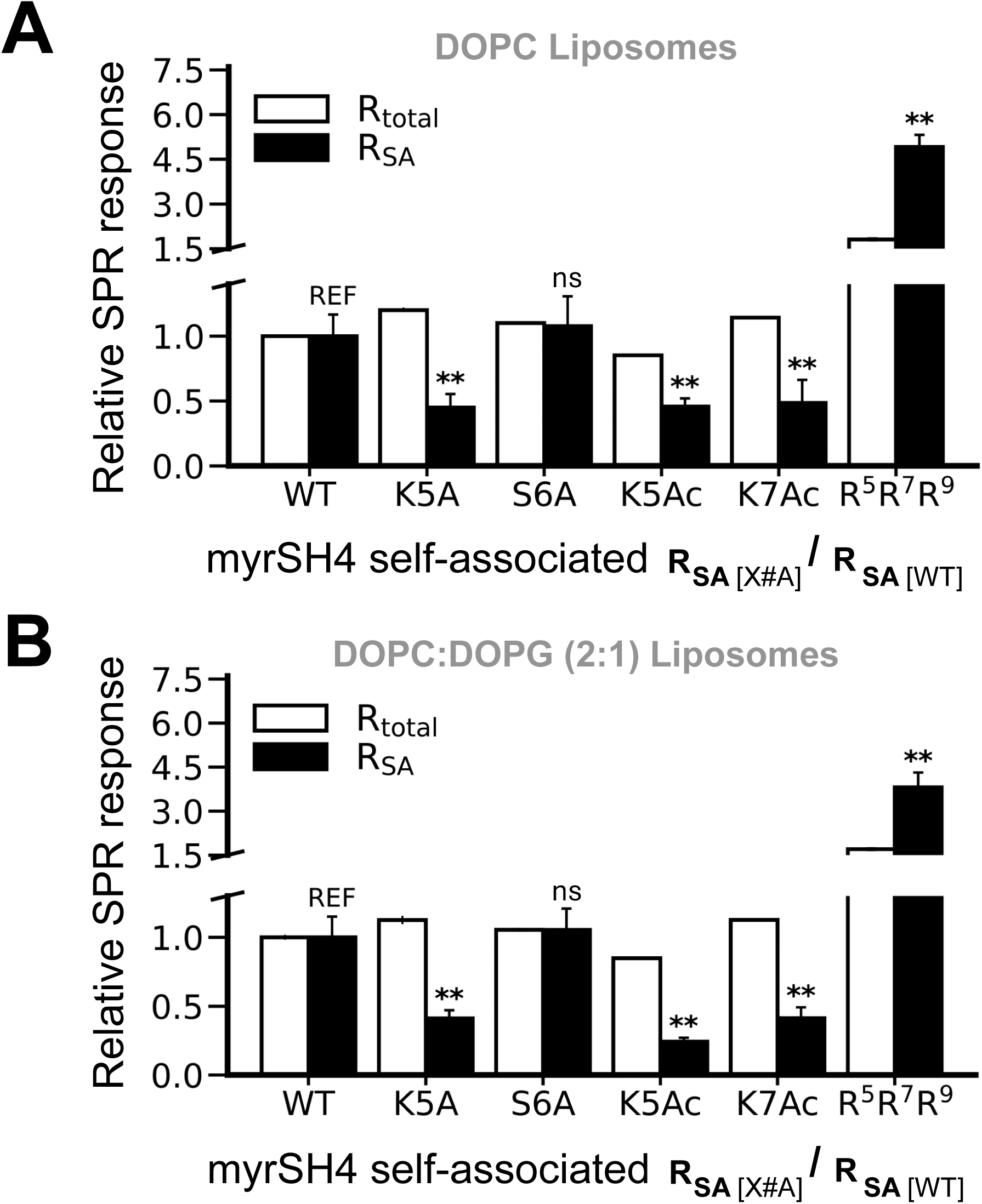
Mutations of lysine residues have similar effects on myrSH4 and SNRE. The effect of the K5A mutation and the lack of effect of the S6A mutation parallel those observed in intact SNRE and presented in Figure 1. The K5A and K7A mutants have a stronger effect on self-association than in the total binding, that is dominated by the monomeric form. The R3 myrSH4 mutant shows a large increase in relative self-association both in the presence of zwiterioninc and negatively charged lipids, emphasizing the fact that a negatively charged phosphate group, to which arginine has a large affinity, is present in both phospholipids.

**Figure S2.**
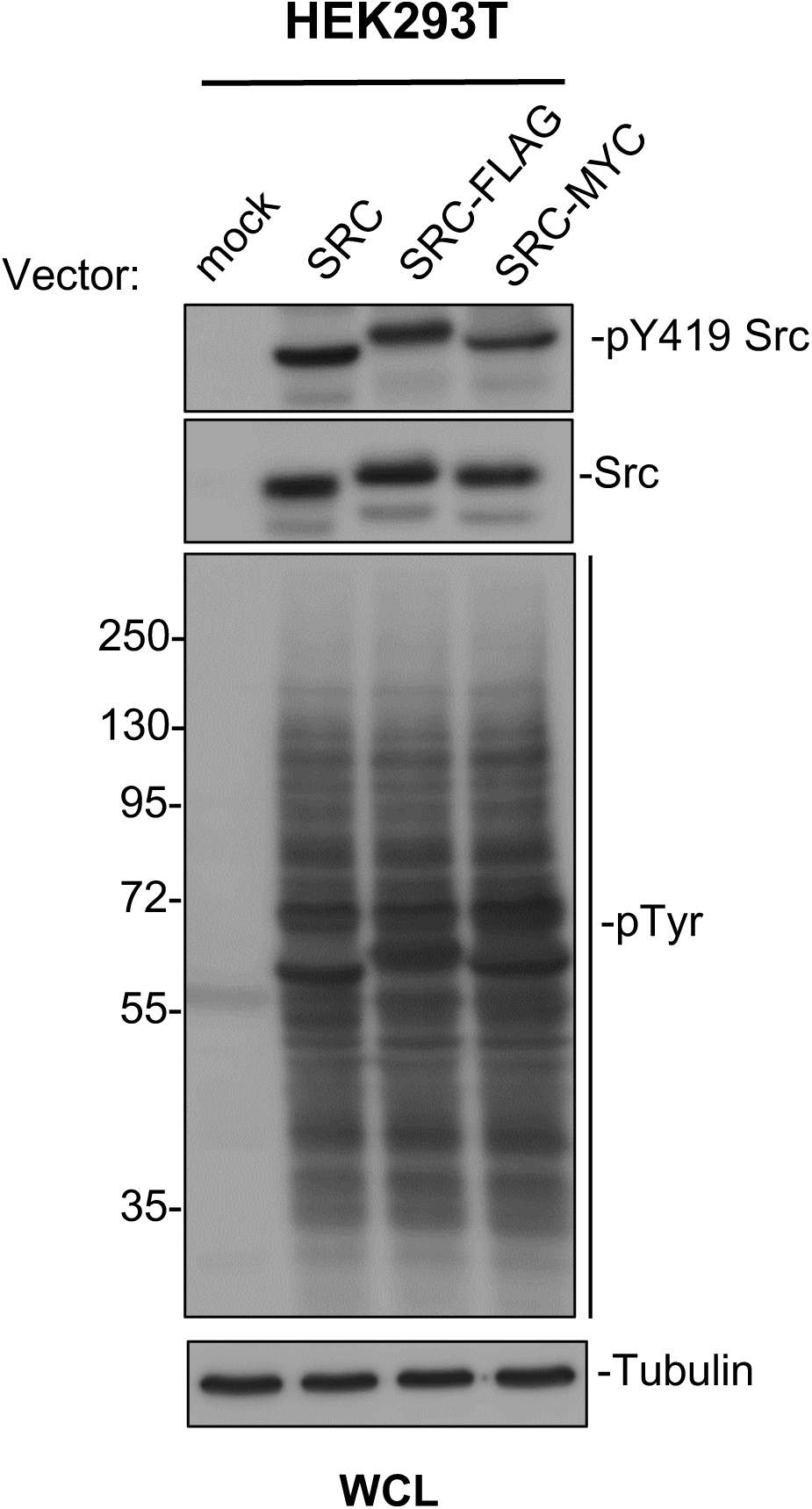
The Flag and Myc tags have no effect on the expression and kinase activity of Src variants. Western blots of whole cell lysates of HEK293T cells retrovirally transfected with WT Src and C-terminal FLAG and Myc tagged variants show similar levels of expression, similar self-phosphorylation, and similar levels of tyrosine phosphorylation.

**Figure S3.**
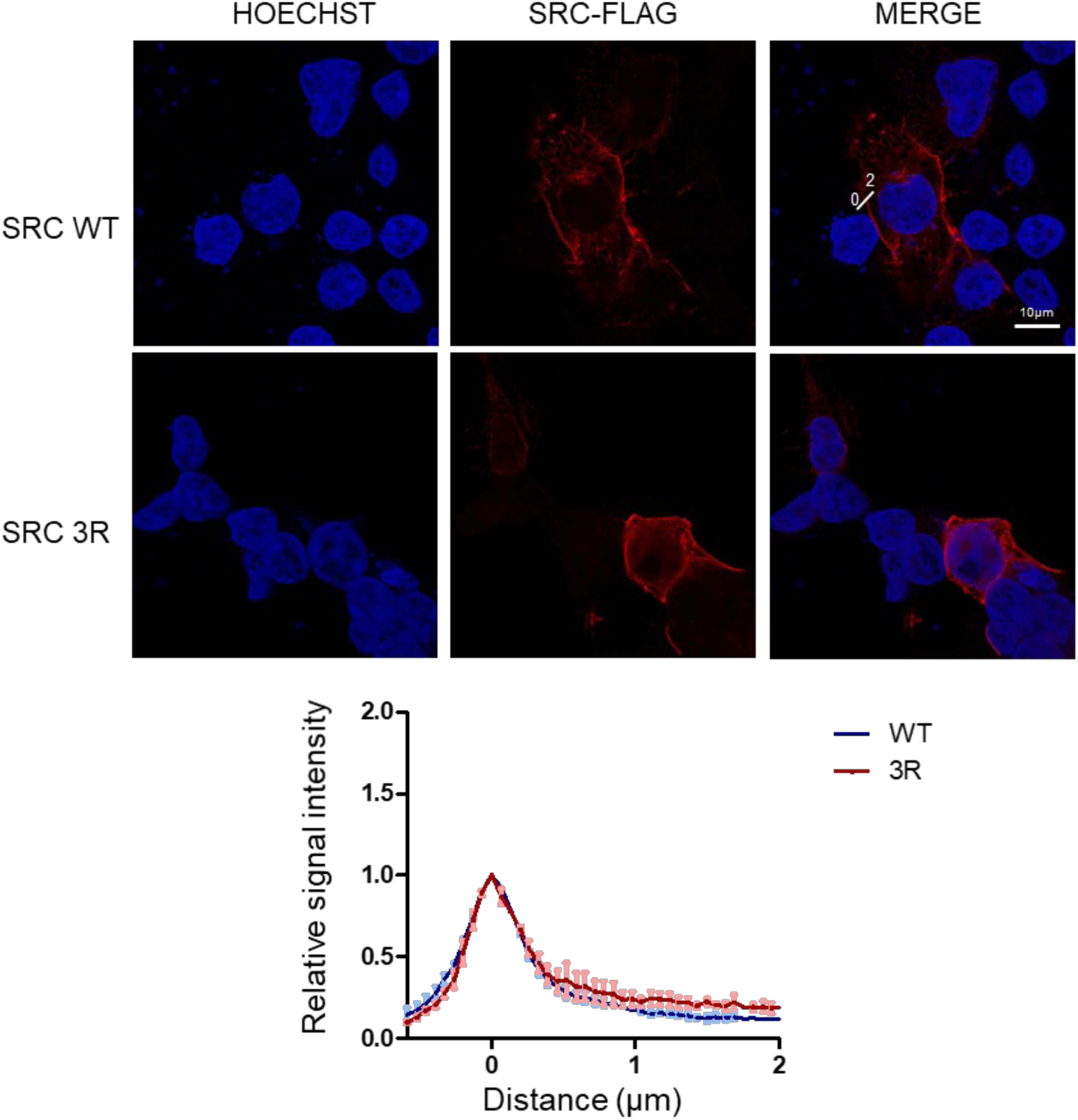
Src WT and 3R colocalize in HEK293T cells. Similar subcellular distribution between Src and Src3R as assessed by confocal microscopy. Subcellular localization of indicated transfected Src Flag construct in HEK293T cells assessed by confocal microscopy of Flag immunofluorescence. Top panel: representative images; bottom panel: quantification (10-12 cells; n=3) relative to the membrane intensity obtained at the plasma membrane for each Src construct.

**Figure S4.**
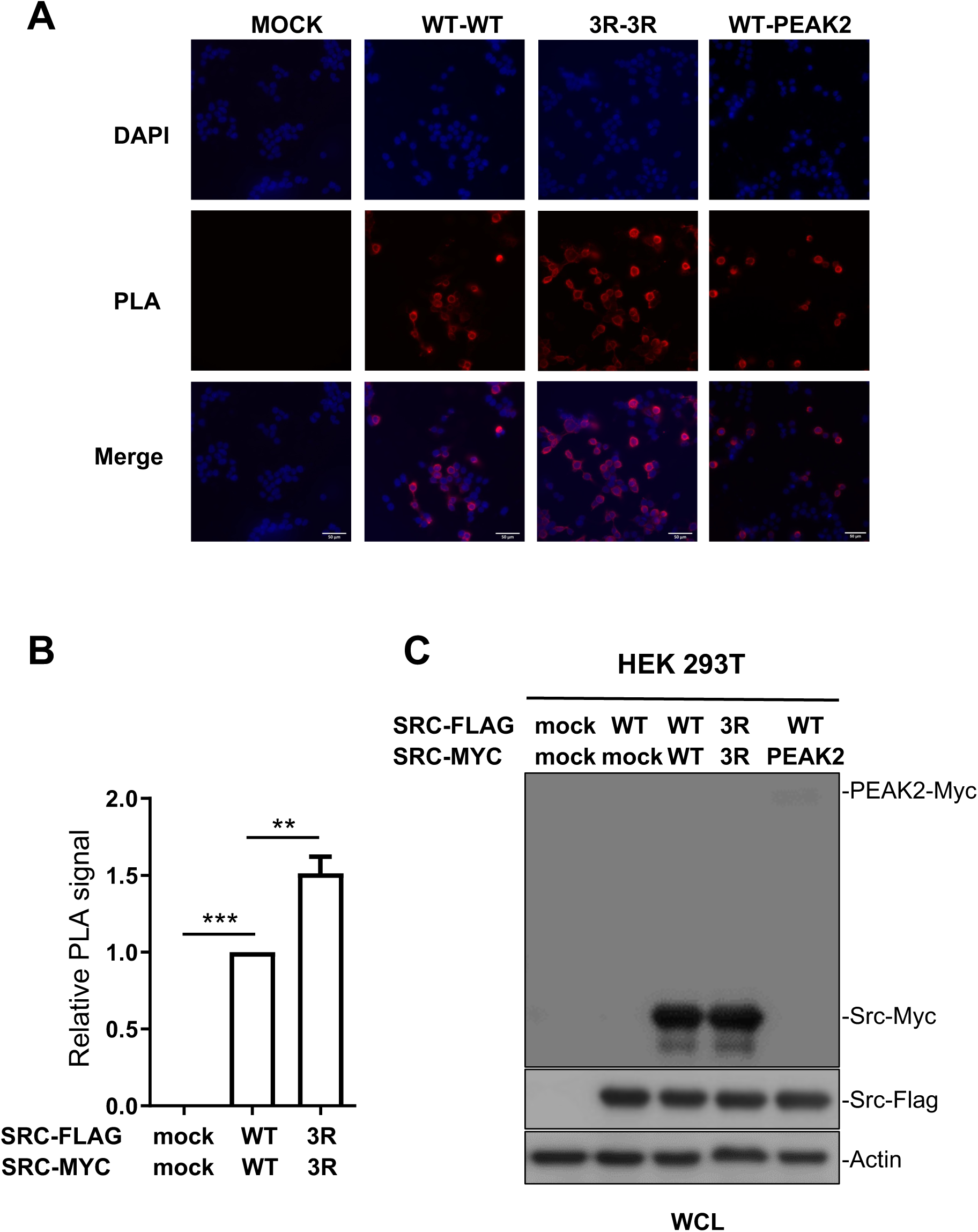
PLA confirms enhanced self-association of the Src 3R mutants. Representative images and quantification by total cell microscopy, showing increased self-association of the Src 3R mutant. A positive control is provided by the known hetero-association of Src with the pseudo-kinase PEAK2. Whole cell lysates confirm that the expression levels of WT and 3R Src variants in very similar. Is shown the mean ± SEM; n=3; **p<0.01; ***p<0.001; Student’s t test

## Material and methods

### Reagents

pFlagN1L11-hSrc and pMycN1L11-Src vectors were described in Aponte et al.^18^ pFlagN1L11-hSrc -Src-K5R/K7R/K9R (Src 3R) was obtained by mutagenesis using the QuickChange Site-Directed Mutagenesis Kit (Stratagene) using specific oligonucleotides as follows: 5’:CCCTTCACCATGGGTAGCAACAGGAGCAGGCCCAGGGATGCCAGCCAGCGG CGCCGC, and 5’:GCGGCGCCGCTGGCTGGCATCCCTGGGCCTGCTCCTGTTGCTACCCATGGT GAAGGG pMX-pS-CESAR retroviral vector expressing human SRC was described in Naudin et al.^28^ pMX-SRC 3R was obtained by inserting BglII-MluI blunt-ended fragments (1687 pb) from pFlagN1L11-hSrc -Src 3R into pMX-pS-CESAR opened by XhoI and blunt-ended.

Antibodies:, anti-Src pY419 (#2101L), anti-Src (clone 327 #ab16885, Abcam) anti-Myc (#2276S) and anti-pTyr clone pY1000 Sepharose bead conjugated (PTM Scan) were from CST, anti-FLAG (M2 antibody, Sigma Aldrich), anti-tubulin (gift from N. Morin, CRBM, Montpellier, France), anti-pTyr 4G10 (gift from P. Mangeat, CRBM, Montpellier, France), anti-cst1 (that recognizes Src, Fyn and Yes) was described in reference ^28^. Anti-rabbit IgG-HRP and anti-mouse IgG-HRP (GE Healthcare).

### Myristoylated proteins expression and purification

Myristoylated SNRE and mutated versions were obtained by co-expression of the N-myristoyl transferase (NMT) enzyme along the Src USH3-His_6_Tag construct in a pETDuet-1 (Novagen) plasmid. The mutations were introduced using QuickChange II XL Site Directed Mutagenesis Kit (Agilent) and Q5 High-Fidelity PCR Kit (New England BioLabs).

Plasmids were transformed in *E. coli* Rosetta^TM^ (DE3)pLysS (Novagen). Bacteria cells were grown in LB medium supplemented with chloramphenicol (25 µg/mL) and ampicillin (100 µg/mL) at 37°C until an OD_600nm_ of ∼0.6. A freshly prepared solution of myristic and palmitic acid (Sigma) (200 µM final concentration/each) and fatty acid free BSA (Sigma) (600 µM final concentration) was added to the cell culture. The lipid-BSA solution was prepared by adding one equivalent of NaOH, heating at 65°C until complete lipid dissolution and adjusting the final pH to 8. Protein expression was induced with 1mM of IPTG (Nzytech) and performed for 5 h at 28°C. 1h after induction start, 6 g/L of glucose was added to the cell culture. Cells were harvested at 4000 rpm for 20 min. Bacteria was lysed in buffer containing 20 mM Tris·HCl, 300 mM NaCl, 10 mM Imidazole, pH 8, 1 % Triton X100 (Sigma), 1x Protein Inhibitor Cocktail (Sigma), 1 mM PMSF (Sigma), 25μg/mL lysozyme (Sigma) and 5μg/mL DNAseI (Roche). Cells were sonicated on ice and centrifuged at 25000 rpm for 45 min. Ni-NTA affinity chromatography was performed using a 1mL Ni-NTA cartridge (GE Healthcare). The protein was eluted with buffer containing 20 mM Tris·HCl, 300 mM NaCl, 10 mM Imidazole, 400 mM imidazole, 0.02 % Triton X100 (Sigma) at pH 8. The final purification step consisted of a size exclusion chromatography in a Superdex 75 26/60 (GE Healthcare) using phosphate buffer (50 mM NaH_2_PO_4_/Na_2_HPO_4_, 150 mM NaCl, 0.2 mM EDTA, pH 7.5). The protein was concentrated with a Vivaspin 20, 5 kDa MWCO concentrator (Sigma Aldrich). The purity of the protein was established by HPLC in a BioSuite pPhenyl 1000RPC 2.0 x 75 mm; 10 μm column coupled to mass spectrometry (ESI-MS), confirming the complete protein myristoylation. All proteins were flash-frozen and stored at -80°C. Myristoylated SH4 domain peptides (myrSH4) were synthesized by SynPeptide Co., Ltd (Shangai, China).

For the AFM experiments, c-Src constructs were myristoylated following an *in vitro* myristoylation with recombinant NMT produced in *E. coli*. SNRE-3R, myrSH4-UD-SH3-SH2, were expressed in a pET28-b vector and full length Src was expressed in a pACYDuet1 together with the Cdc37 chaperone in cells coexpressing GST-tagged YopH tyrosine phosphatase. All constructs were produced as N-terminal His-tagged SUMO fusion, that was used for affinity purification and removed by Ulp1 treatment to provide the native N-terminal sequence ready for myristoylation. All synthetic genes were acquired from the GenScript.

Plasmids were transformed in *E. coli* Rosetta^TM^ (DE3)pLysS (Novagen). Bacteria cells were grown in LB medium supplemented with Kanamycin (50 µg/mL) and chloramphenicol (25 µg/mL) at 37 °C until an OD_600nm_ 0.6-0.8 was reached. Expression was induced with 0.5 mM IPTG and left overnight at 30 °C. Cells were harvested at 4000 rpm for 20 min. Bacteria were resuspended in buffer containing 50 mM NaH_2_PO_4_/Na_2_HPO_4_, 300 mM NaCl, 20 mM Imidazole, pH 8, 1 mM PMSF (Sigma), 1 mM Benzamidine (Sigma), 25μg/mL lysozyme (Sigma) and 5μg/mL DNaseI (Roche). Cells were sonicated on ice and centrifuged at 25000 rpm for 30 min. Ni-NTA affinity chromatography was performed using a 5 mL Ni-NTA cartridge (GE Healthcare). The protein was eluted with buffer containing 50 mM NaH_2_PO_4_/Na_2_HPO_4_, 300 mM NaCl, 400 mM imidazole, pH 8. The eluted sample was buffer exchanged using size exclusion chromatography in a Superdex 75 26/60 (GE Healthcare) to a myristoylation buffer (50 mM NaH_2_PO_4_/Na_2_HPO_4_, 100 mM NaCl, pH 7.3).

Full length Src was expressed in E.coli B834(DE3) competent cells. Purification included Ni-NTA affinity purification, Ulp1 cleavage, GSTrap HP (Cytiva) to remove residual GST-tagged YopH that coeluted in the previous purification steps, HiTrap Q HP ion exchange, and a final purification with a Superdex 75 26/60 size exclusion column.

For in vitro myristoylation, constructs and myristoyl coenzyme A (M4414, Sigma-Aldrich) were mixed at a 1:2 ratio. The mixture was supplemented with 2 μM of his-tagged recombinant human-NMT,1 mM DTT, and 30 mM CHAPS and left at room temperature for 2 hours. Then, the sample was injected into a 1 mL HiTrap Q HP column to remove the detergent and eluted with a salt gradient. NMT was removed with Ni-NTA beads. The myristoylated constructs were buffer exchanged to PBS (137 mM NaCl, 2.7 mM KCl, 10 mM Na_2_HPO_4_, and 1.8 mM KH_2_PO_4_) using a Superdex 75 26/60 (GE Healthcare). The myristoylation state of the protein was established by HPLC in a BioSuite pPhenyl 1000RPC 2.0 x 75 mm; 10 μm column coupled to mass spectrometry (ESI-MS).The protein was concentrated with a Amicon Ultra - 15, 10 kDa MWCO concentrator (Merck Millipore). All proteins were flash-frozen and stored at - 80°C.

### Preparation of liposomes

1,2-dioleoyl-sn-glycero-3-phosphocoline (DOPC) (TebuBio) and 1,2-dioleoyl-sn-glycero-3-phospho(1’-rac-glycerol) (sodium salt) (DOPG) (Sigma) were dissolved in chloroform. Lipid compositions used were DOPC and DOPC:DOPG (2:1 or 4:1) at 4mM. The organic solvent was evaporated under a nitrogen stream. Lipid films were rehydrated with vortexing in phosphate buffer 50 mM NaH_2_PO_4_/Na_2_HPO_4_, 150 mM NaCl, 0.2 mM EDTA, pH 7.5 (SPR assays) or 10 mM NaH_2_PO_4_/Na_2_HPO_4_, 150 mM NaCl, 5 mM MgSO_4_, pH 7.5 (AFM experiments). Large unilamellar vesicles were prepared by extrusion using a Mini-Extruder (Avanti Polar Lipids). The lipid suspension was extruded 15 times through a 100 nm polycarbonate filter. The mean diameter of the liposomes was verified by Dynamic Light Scattering (Zetasizer Nanoseries S. Malvern instruments). Liposomes were used within two days to avoid lipid oxidation.

### Surface Plasmon Resonance assays

The complete protocol (Figure S1) consists of: 1) liposome capture, 2) myristoylated protein binding (including reversible and irreversible bound forms) 3) washing of the major monomeric fraction, 4) antibody binding to the remaining, minor persistently bound fraction and 5) regeneration with detergents to strip off the captured liposomes. The formation of the lipid bilayer and primary c-Src binding produced clear SPR responses. Persistently bound c-Src dimers represent a minor fraction of the steady-state bound protein causing only a weak shift from the baseline (less than 5% of the primary response dominated by binding of c-Src monomers), thus the response is amplified with the antibody capture, so that the relative amount of dimer formed by c-Src variants can be accurately quantified. We used an antibody directed to the His_6_Tag located at the C-terminus to minimize the interference with the membrane-anchoring region. For myristoylated peptides, lacking a His_6_Tag we used an antibody directed to the Src-N-terminal region. The corresponding mutations in synthetic peptides and longer Src constructs showed equivalent effects, confirming the antibody against Src N-terminus was not inducing any artifact.

All measurements were carried out in a Biacore^TM^ T200 instrument (Cytiva) at 25°C. A 2D-carboxylmethyldextran sensor chip (Xantec) was used. All the flow cells, except for the reference, were modified by a covalent attachment of phytosphingosine (TebuBio) to allow the capture of liposomes. An amine-coupling procedure was performed with 1 mM of phytosphingosine in freshly prepared 20 mM sodium acetate buffer at pH 6.7. The running buffer for all experiments consisted of 50 mM NaH_2_PO_4_/Na_2_HPO_4_, 150 mM NaCl, 0.2 mM EDTA, pH 7.5. DOPC and DOPC:DOPG (2:1) liposomes at 1 mM concentration were coated over the phytosphingosine containing flow cells by a 20 s injection at 10 µL/min. In case the distinct flow cells contained different types of lipids, the DOPC liposomes were injected in the first available flow cell to avoid anionic lipid migration towards the neutral liposomes through the flow cells. The reference cell and possible uncovered surface in the liposomes flow cells were blocked with 1 mg/ml of BSA at 50 µL/min for 20 s. Mass transport effects were minimized by injecting the myristoylated c-Src variants at 50 µL/min. Protein concentration ranged from 3 µM to 20 µM for the protein density experiments and it was fixed to 20 µM for the mutants analysis. The myrSH4 peptides were injected at 50 µM. The protein was allowed to associate for 60 s while the dissociation lasted 350 s. The antibody (antiHis_6_Tag or antiSrc antibodies (both Santa Cruz Biotechnology) was injected at 1:5 dilution in running buffer for 60 s at 30 µL/min. n≥3 experiments were performed for the c-Src variants, each time in randomized order. The surface was regenerated with two pulses (30 s at 100 µL/min) of Isopropanol:50 mM NaOH (2:3) solution followed by a 20 mM CHAPS (Sigma) or 40mM Octyl-beta-Glucoside (Sigma) pulse. Each cycle started with freshly captured liposomes. All data was double referenced (reference channel and baseline subtraction (0μM concentration) and analyzed using BiaEval 3.1 software (GE Healthcare). Experiments were performed in two sensor chips, reproducible as indicated in Fig. 2C.

### Data and statistical analysis

The different mutants were compared relatively to WT using the following equation.

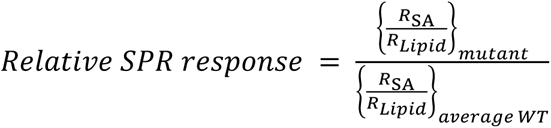

Where R is the SPR response observed in the corresponding sensorgram. All the statistical analysis were performed using the Python scientific library SciPy (SciPy: Open-Source Scientific Tools for Python, http://www.scipy.org, last accessed Apr 12, 2022). If not otherwise stated, the analysis was performed against the WT c-Src construct. Unpaired two-sample t-test (two-sided) was conducted.

### AFM Experiments

SLBs were obtained by direct fusion of liposomes onto freshly cleaved mica surfaces (mica discs, Ted Pella, Redding, CA). 80 µL of DOPC or DOPC:DOPG (4:1) liposomes (2 mM concentration in 10 mM NaH_2_PO_4_/Na_2_HPO_4_, 150 mM NaCl, 5 mM MgSO_4_, pH 7.5) were placed onto the mica surface. After 30 min at room temperature, the samples were rinsed several times with working buffer (10 mM NaH_2_PO_4_/Na_2_HPO_4_, 150 mM NaCl, pH 7.5) to remove unfused vesicles, keeping the membranes always hydrated. Atomic force microscopy (AFM) imaging was performed with a NanoWizard 3 BioScience AFM (JPK Instruments, Bruker Nano GmbH) at room temperature and under liquid environment (working buffer). Silicon nitride probes with a nominal spring constant of 0.35 N/m (DNP-S10, Bruker) were used. After having measured the sensitivity (V/m), the cantilever spring constants were individually calibrated using the equipartition theorem (thermal noise routine). Imaging was performed in AC mode. Lipid membrane formation was verified by imaging and performing force-distance curves on different areas of the samples. Then, protein (or myrSH4 peptide) was added to the buffer solution covering the sample, to a final concentration of 100 nM (or 600 nM for the myrSH4 peptide) and imaging was restarted after 4 min.

### Cell transfection and retroviral infection

(HEK293T and HCT116 cells) (ATCC, Rockville, MD) were cultured in Dulbecco’s Modified Eagle’s Glutamax Medium (DMEM, Invitrogen) supplemented with 10% fetal bovine serum, 100U/mL penicillin and 100μg/mL streptomycin. Transfections and retroviral infections were done as described in Aponte et al. ^18^

### Proximity Ligation Assay and confocal microscopy

HEK293T cells plated on glass coverslips were transfected with Src-Flag and Src-Myc mutants constructs for 48 hours and subcellular SRC distribution was analyzed after cell fixation (4% paraformaldehyde, 0.5% triton, for 20min) by fluorescence using fluorescent microscopy as described in Aponte et al. ^18^ Proximity Ligation Assay (PLA) was performed according to the manufacturing kit protocol (#DUO92101, MERCK). Briefly, HEK293T cells plated on glass coverslips were transfected with Src-Flag and Src-Myc mutants or PEAK2-Myc (as a positive control)^27^ constructs for 48 hours and fixed with 4% paraformaldehyde, 0.5% triton during 20 min. Glass slides was blocked in the Duolink® Blocking Solution during 1 hour at 37°C and then incubated in primary antibodies rabbit anti-Flag (#F7425, Sigma, dilution 1/250) and mouse anti-Myc (#9B11, Cell Signaling, dilution 1/500) in the Duolink® Antibody Diluent during 1 hour at room temperature. After the washing steps, the slides were incubated with the PLUS and MINUS PLA diluted in the Duolink® Antibody Diluent (1:5) during 1 hour at 37°C. After three washes, the slides were incubated in the ligation solution during 30 min at 37°C and the amplification solution. Finally, after washes the slides were mounted with the Duolink® In Situ Mounting Medium with DAPI and observed with an upright fluorescent microscope. Confocal microscopy analysis was described in Aponte et al.^18^ Briefly, images were acquired using a Zeiss LSM980 NLC confocal microscope equipped with a ×63 plan-apochromat oil immersion objective (NA 1.4) and a pinhole set to 1 Airy, controlled by Zen acquisition software (bleu edition). Optical sectioning was set to 1 μm and digital images were further processed using Fiji software and the intensity line plot analysis plug-in (https://fiji.sc/). Line profiles were extracted as a text file and calculated in Excel to manually detect the cell boundary, which was set to 0 and used as the intensity normalisation point, then normalised data were collected and graphically and statistically analyzed using Prism software. (10-15 images per biological replicate, n=3). Functional assays in cells. Soft agar and cell migration assays were described previously^28,18^ For colonies formation assay in soft agar, 1 500 cells per well were seeded in 12-well plates in 1ml DMEM containing 10% FCS and 0.33% agar on a layer of 1ml of the same medium containing 0.7% agar. After 18-21 days, colonies with > 50 cells were scored as positive. Cell migration assay was performed as described in (32) using Fluoroblok invasion chambers (BD Bioscience). 5 000 cells per well on top in 0% FCS medium and 10% FCS medium at the bottom. After 30 min at 37°C, the cells were labelled with calcein AM (8µg/ml, Sigma-Aldrich) and migrative cells were photographed using the EVOS FL Cell Imaging System. Quantification of the number of invasive cells per well was done with the ImageJ software.

